# Genetic Mapping in Diversity Outbred Mice Identifies Novel Loci and Candidate Genes for Anxiety-Like Behavior and Genetic Subgroups Predictive of Ethanol Consumption

**DOI:** 10.1101/2025.05.27.656445

**Authors:** Zachary Tatom, Kristin M. Mignogna, Lorna Macleod, Zachary Sergi, Michael F. Miles

## Abstract

Anxiety disorders are the most common class of psychiatric disorder. Risk for anxiety disorders is thought to be influenced by many genes, each contributing a small effect. The light-dark box behavioral assay was designed to measure anxiety-like behavior in rodents. Diversity Outbred (DO) mice were designed for high-resolution quantitative trait loci (QTL) mapping on a genetically-diverse background. Here, we studied a population of 518 male DO mice for anxiety-like and locomotor behaviors from a light-dark box assay. Multivariate analysis of behavioral data identified two major subgroups of animals differing in basal anxiety behavior and subsequent ethanol consumption patterns. Behavioral QTL analysis identified a significant locus on Chromosome 14 associated with 3 anxiety-like behavioral phenotypes. Haplotype analysis revealed an effect of C57BL/6J alleles at this locus, with mice carrying those alleles exhibiting more anxiety-like behavior. An additional 9 suggestive loci were identified. Genes located within the confidence intervals for the Chromosome 14 locus were analyzed for coding sequence polymorphisms, prefrontal cortex expression QTLs, human GWAS data, and additional data sets related to psychiatric conditions including substance use. Results prioritized two candidate genes, *Tbc1d4* and *Lmo7*, for further study. These results represent the highest-resolution genetic mapping of light-dark box behaviors in mice to date, revealing insights into the complex biology of anxiety disorders. Additionally the studies identify striking subgroups of animals where basal anxiety-like behavior predicts eventual ethanol consumption phenotypes.

## Introduction

Anxiety disorders are a major public health concern affecting an estimated 374 million people worldwide in 2020, making them the most common class of mental health disorder [1, 2]. In the United States it has been estimated that 31.1% of adults are expected to experience any anxiety disorder in their lifetime [3, 4]. Risk for anxiety disorders is influenced by complex genetic and environmental factors; a 2015 meta-analysis of over 2,500 twin studies estimated overall anxiety disorder heritability at about 49% [5], while more recent heritability estimates from GWAS are around 26% [6], though estimates differ by individual anxiety disorder. Anxiety disorders are also highly comorbid with other mental health disorders including alcohol use disorder (AUD), with individuals diagnosed with AUD having 2.1 times greater risk of having any anxiety disorder compared to non-alcohol users (OR 2.1, 95% CI 2.0-2.2) [7]. Numerous human GWAS studies have identified loci associated with anxiety disorders, implicating genes including CTNND1, RAB27B [8], SATB1, MAD1L1 [9], and others. However, the individual genes associated with anxiety disorders are thought to be of small effect [10, 11], with recent estimates indicating as many as 12,900 potential risk loci [12]. Moreover, characterizing the biological mechanisms driving associations between identified risk variants and anxiety disorders also remains a challenge thus warranting further study to identify risk loci and characterize their effects.

Additionally, heterogeneity in anxiety and environmental contributions to the trait complicate human studies [13]. In preclinical research, the light-dark box (LDB) assay is widely used in rodent models to assess unconditioned anxiety-like behaviors [14–16]. The assay is thought to take advantage of the interplay between a rodent’s natural exploratory drive and its aversion to brightly-lit, open spaces [17, 18]. Transitions between light and dark chambers, total locomotor activity, time spent in the light chamber, and distance traveled in the light chamber have all been used as quantitative measures of anxiety-like behavior [16, 18–21]. Inbred crosses have been used to create genetic maps of these behaviors, with relatively large confidence intervals [22–24] which make identifying individual candidate genes associated with phenotypes difficult.

Recent development of mouse genetic models with high genetic diversity and recombination have greatly improved sensitivity in identifying quantitative trait loci (QTL) [25–28]. Diversity Outbred (DO) mice, derived from outbreeding of 8 genetically- and phenotypically-diverse founder strains [29], provide high levels of allelic variation and recombination events enabling high-resolution genetic mapping [26].

Our lab has previously used a population of nearly 600 Diversity Outbred mice to map QTLs associated with voluntary ethanol consumption behaviors at a higher resolution than previously seen [30, 31]. Here we describe the first use of DO mice to map behavioral QTLs associated with anxiety-like behaviors in the light-dark box, prioritizing candidate genes based on haplotype analysis and bioinformatics studies.

## Methods

### Ethics Statement

All animal care and euthanasia procedures were performed in accordance with the rules and regulations established by the United States Department of Agriculture Animal Welfare Act and Regulations, Public Health Services Policy on Humane Care and Use of Laboratory Animals, and American Association for Accreditation of Laboratory Animal Care. Humane endpoints were established by the same standards.

### Animal Studies

Male DO mice (*n* = 636) were acquired from Jackson Laboratories after weaning at 4-6 weeks of age in 7 cohorts (average *n* = 106) spanning DO generations 22-25. Mice were singly housed in temperature- and humidity-controlled vivariums on cedar shaving bedding with *ad libitum* access to water and standard chow (#7912, Harlan Teklad, Madison, WI, United States).Vivariums were set up with alternating 12-hour light and dark phases, and all mice were weighed weekly. Experimental procedures were begun on animals at 8-12 weeks of age. Sample size considerations and exclusion of female mice were chosen to maximize power to detect significant QTL. Study of behaviors in ∼600 animals was predicted to have 80% power to detect alleles causing less than 5% variance in a trait at a p-value of <0.05 [32].

### Behavioral Phenotyping via Light-Dark Box

In order to assay locomotor activity and anxiety-like behavior, a Med-Associates Inc. (Fairfax, VA) Mouse Locomotor Box with light-dark inserts was used. Animals were acclimated to the behavioral testing room for 1 hour prior to behavioral testing. Mice were placed in the box in the transition area between light and dark sides facing the dark half and their movements were tracked via laser beams between the sides of the box for 10 minutes. Total locomotor activity and total jump counts were measured across this time period, along with the number of transitions between light and dark chambers and distance traveled and time spent in either the light or dark chamber. Percent distance traveled, percent time spent, and percentage of jumps in the light chamber were derived as additional anxiety-like behavioral phenotypes, with animals that travel more in the light or spend more time in the light considered to exhibit lower levels of anxiety-like behavior.

### Behavioral Phenotyping via Intermittent Ethanol Access

Beginning 24 hours after LDB, animals were exposed to 5 weeks of voluntary ethanol consumption using a 3-bottle choice intermittent ethanol access (IEA) procedure described in detail elsewhere [33]. Briefly, mice were exposed in alternating 24-hour periods at the beginning of the dark cycle to three bottles (H_2_O, 15% v/v EtOH, 30% v/v EtOH) and consumption was measured at the end of each 24-hour period. Total ethanol consumption in g EtOH per kg mouse body weight, ethanol preference as percentage of ethanol consumed compared to total fluid consumed, and preference for 30% v/v EtOH compared to total ethanol consumed were calculated and used for comparison with light-dark box phenotypes. Only the first 4 weeks of IEA were used for these analyses.

### Tissue Sample Collection and Genotyping

Mice were euthanized 24 hours after the end of their last ethanol exposure period via cervical dislocation and decapitation and tissue samples were collected immediately afterwards, flash-frozen in liquid nitrogen and stored at −80° C. Brains were microdissected into nine regions as previously described [34, 35]. Tail snips were collected for genotyping (NeoGen Inc; Lincoln, NE) using a GigaMUGA microarray (*n*_SNP_ = 141,090; *n*_CNV_ = 2,169) designed to optimize genotyping of DO mice.[36] Of the initial 636 DO mice, full datasets were only collected from 630 mice as 6 mice reached humane endpoints before the end of experimentation. Quality control removed an additional 29 mice (22 for poor genotyping quality and 7 for potential sample mix-ups), and an additional 110 mice were removed from light-dark box studies resulting in the final sample of 493 used for analyses presented here.

### RNA-seq from Prefrontal Cortex

Prefrontal cortex (PFC) samples were taken from 220 of the Diversity Outbred mice 24 hours after the end of IEA. 20 of these mice were ethanol-naïve controls, whereas the other 200 were chosen based on week four of IEA to identify the 100 highest-drinking mice and 100 lowest-drinking mice. RNA-seq methods are described in another manuscript [30, 33].

### Behavioral Data Analysis

Phenotypic data from LDB were square-root transformed for normalization, whereas ethanol consumption data were log transformed and ethanol preference data were square-root transformed. Heritability estimates were generated using R/qtl2 software for R [37]. A kinship matrix was estimated using the “overall” method in R/qtl2. Heritability of each light-dark box phenotype was estimated using restricted maximum likelihood and kinship estimates.

Because LDB occurred prior to the initiation of IEA, anxiety-like phenotypes were used as predictors in linear models (using ordinary least squares regression) to estimate mean ethanol consumption over four weeks of IEA. Furthermore, *k*-means clustering was used to assign mice to one of two clusters based on anxiety-like behavioral data [38]. Mean ethanol drinking behaviors were examined across clusters using *t*-tests. In order to determine if cluster membership had any effect on the trajectory of ethanol consumption over time, a linear model was constructed to regress consumption by drinking day, cluster membership, and an interaction term between the two predictors.

### QTL Analysis

Significant covariates were identified using linear modeling in R to regress each behavioral phenotype on each potential covariate. Behavioral QTL (bQTL) mapping was then carried out using R/qtl2. Kinship between mice was included as a covariate in all QTL analyses and was estimated using a linear mixed model and the leave-one-chromosome-out (LOCO) method. Cohort and body weight were used as additive covariates for total locomotor activity, transitions between chambers, and percent time spent in the light. For percent distance traveled in the light, only cohort was used because body weight was not a significant covariate for that phenotype. For gene expression QTL (eQTL) from PFC, only cohort was used as a covariate. eQTL were filtered to those within 1 Mbp of the gene being transcribed to identify *cis*-eQTL.

Logarithm of the Odds (LOD) scores were calculated for each marker and permutation analysis (*n*_perm_ = 1000) used to calculate genome-wide empirical *p*-values. SNP variant LOD scores were similarly calculated for QTL intervals and significance level *p*-values determined by permutation across the involved chromosome. To detect founder strain effects, haplotype analysis was conducted for chromosomes containing significant or suggestive bQTLs using best linear unbiased predictors (BLUPs) in R/qtl2. SNP associations were also identified using R/qtl2 software and compared to Mouse Genome Informatics (MGI) databases to identify consequences and variant founder strain distribution patterns [39]. Significance thresholds for SNPs were identified using permutation analysis across the chromosome containing a QTL (*n*_perm_ = 1000). Genes were identified within 95% C.I.s using MGI databases.

### Differential Expression Analysis

DESeq2 software for R was used to identify significantly differentially-expressed genes and estimate fold changes across *k*-means clusters using RNA-seq data from both PFC (*n* = 17) and NAc (*n* = 36) tissues from ethanol-naïve mice [40]. A generalized linear model was used and significance thresholds tested using a Wald test to compare gene expression by contrast. Untransformed count data were used, the null hypothesis was set to a log fold change of 0 (indicating no differential expression), and *p*-values were corrected for false discovery rate using the Benjamini-Hochberg method. After analysis, results were filtered to identify differentially-expressed genes as those with both a corrected *p*-value less than or equal to 0.05 and an absolute value log fold change (LFC) greater than 0.1.

### Bioinformatics Analysis

Positional candidate genes from significant bQTL from LDB were searched for in GeneWeaver to find data sets relevant to psychiatric conditions [41]. Additionally, GWAS Catalog was searched to identify associations between human orthologs of positional candidate genes and anxiety-related traits [42].

## Results

### DO Mice Exhibit Phenotypic Diversity in Light-Dark Box Assay

Diversity Outbred mice display phenotypic variance in LDB behaviors (Table 1, Figure 1A). SNP-based heritability estimates ranged from 0.104 for percent jumps in the light to 0.599 for distance traveled (Figure 1B). All LDB behaviors were significantly correlated, with Spearman *rho* values ranging from 0.2 – 0.76 (Figure S1). The variance in phenotype and heritability of each behavior indicates that they are good candidates for QTL analysis.

**Figure 1.**
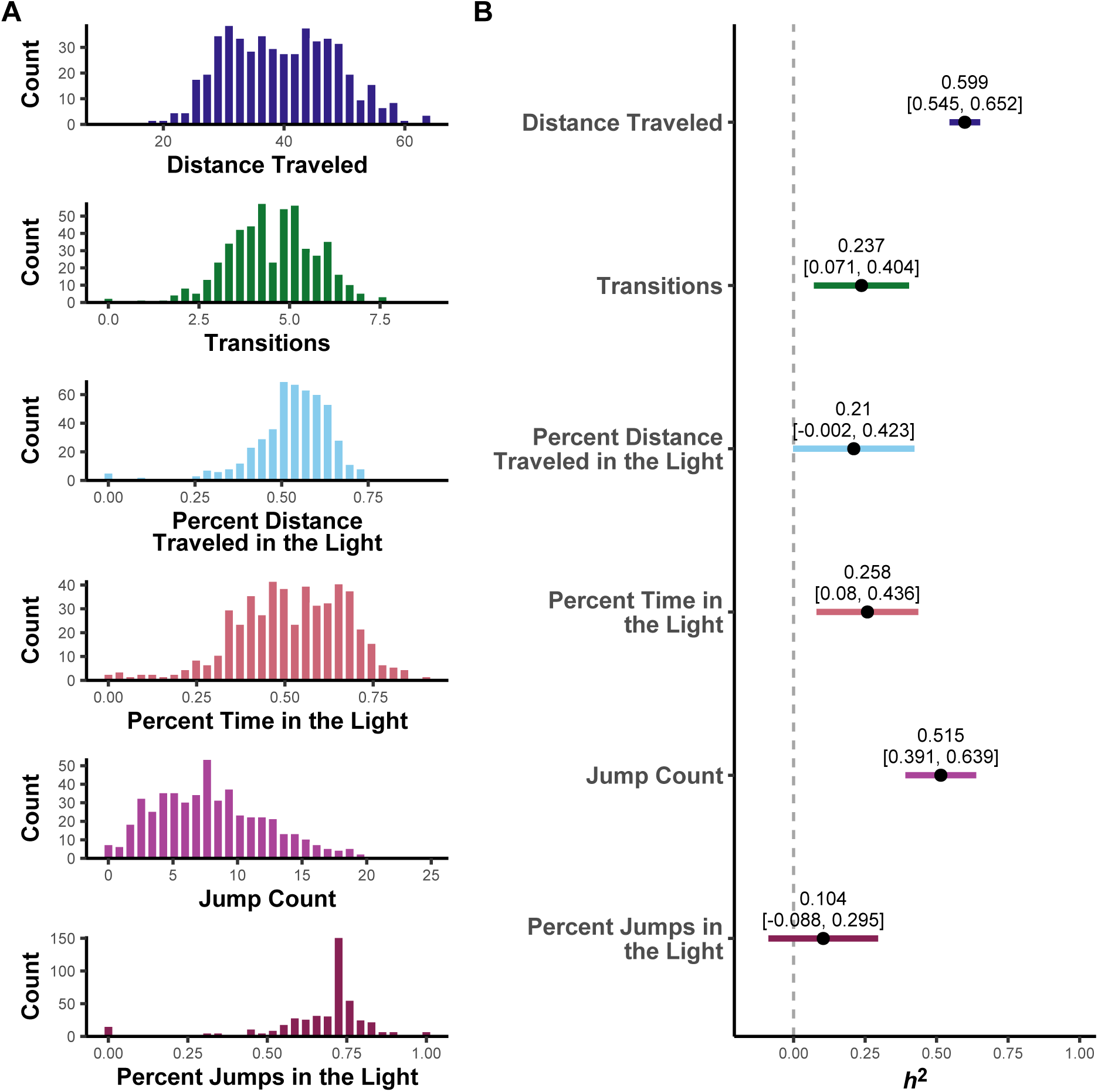
Normalized light-dark box phenotypes and SNP-based heritability estimates. All phenotypes from light-dark box assay were square-root transformed for normalization (A). These normalized phenotypes were then used to estimate heritability in the DO mouse population (B). 95% confidence intervals are presented in brackets. Of the phenotypes analyzed, only percent distance traveled in the light chamber and percent jumps in the light chamber had confidence intervals including zero, indicating that most of these phenotypes are estimated to be significantly heritable in this mouse population.

**Table 1.** Summary statistics from light-dark box phenotypes. Data were collected from Diversity Outbred mice undergoing light-dark box assay over the course of10 minutes. All phenotypes are square root-transformed for normalization.

| **Phenotype** | ***n*** | **Mean** | **Median** | **Std.**<br>**Dev.** | **Min** | **Max** |
| --- | --- | --- | --- | --- | --- | --- |
| Distance Traveled | 518 | 39.49 | 39.19 | 9.38 | 11.24 | 63.91 |
| Transitions | 518 | 4.53 | 4.58 | 1.24 | 0.00 | 8.77 |
| Percent Distance Traveled in the Light | 518 | 0.52 | 0.54 | 0.13 | 0.00 | 0.92 |
| Percent Time in the Light | 518 | 0.52 | 0.52 | 0.16 | 0.00 | 0.90 |
| Jump Count | 518 | 8.10 | 7.55 | 4.51 | 0.00 | 24.62 |
| Percent Jumps in the Light | 518 | 0.66 | 0.71 | 0.18 | 0.00 | 1.00 |

*k*-means clustering identified two potential clusters in the data (Figure S2A). Cluster 1 contained 262 mice and generally correlated with increased locomotor activity, transitions, percent distance traveled in the light, percent time spent in the light, jump count, and percent jumps in the light compared to Cluster 2, which contained 255 mice. This indicates decreased anxiety-like behavior in Cluster 1 compared to Cluster 2.

### Light-Dark Box Phenotypes Predict Later Voluntary Ethanol Consumption in DO Mice

Linear regression models were created to examine relationships between individual LDB phenotypes and voluntary ethanol consumption (Figure 2). Each LDB phenotype was significantly predictive for mean voluntary ethanol consumption across 4 weeks, with all coefficient estimates being positive indicating that increased locomotor activity (Figure 2A) and decreased anxiety-like behavior (Figure 2B-F) predict increased ethanol consumption.

**Figure 2.**
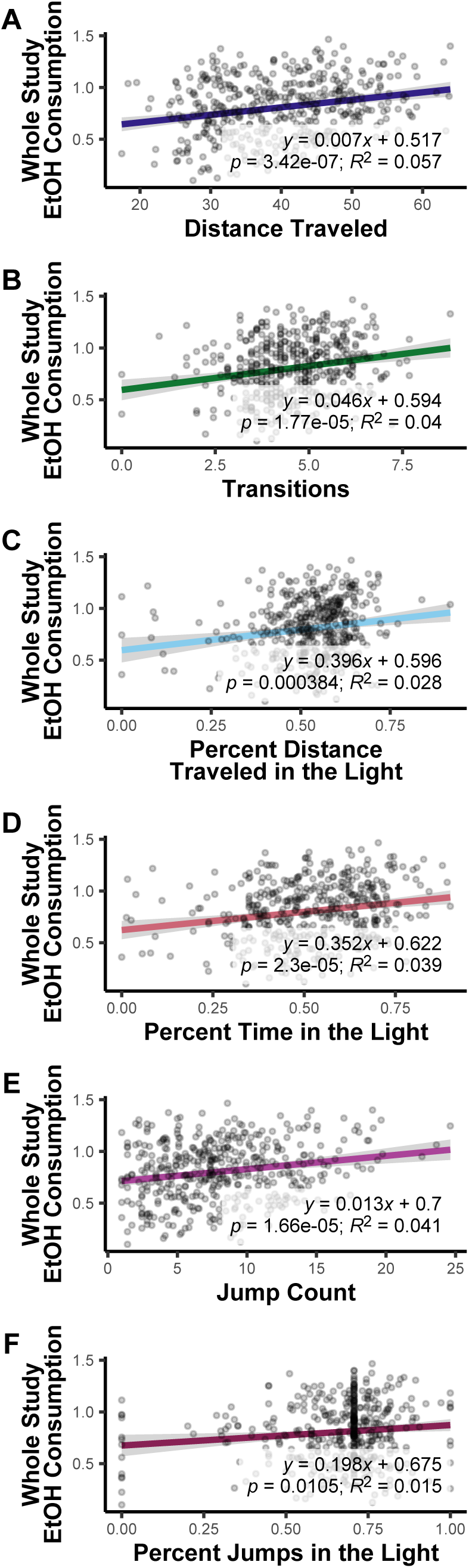
Anxiety-like behaviors significantly predict voluntary ethanol consumption in Diversity Outbred mice. Linear models used individual light-dark box phenotypes to predict mean voluntary ethanol consumption over 4 weeks of intermittent ethanol access. Light-dark box phenotypes were square-root transformed for normalization; ethanol consumption was calculated as g EtOH / kg mouse body weight and log-transformed for normalization. All light-dark box phenotypes were significant predictors for ethanol consumption, explaining between 1.5% for percent jumps in the light (F) and 5.7% for distance traveled (A) of the variance in consumption.

Means for ethanol consumption (Figure S3A), preference for ethanol over water (Figure S3B), and preference for a higher concentration of ethanol (Figure S3C) were compared across *k*-means clusters generated from light-dark box data. Mice in Cluster 1 consumed significantly more ethanol across the study (Figure S4A) and showed increased preference for 30% ethanol; in weeks 1 and 4, they also showed increased preference for ethanol compared to water, but not when averaged over the whole study.

To determine if clusters also differed in the rate at which ethanol consumption over time increased, a linear model was fitted to predict consumption using drinking day, cluster, and an interaction term (Table S1). This model showed a significant difference in slopes between the two clusters (Figure S4B) as indicated by the interaction term, with Cluster 1 mice showing a steeper increase in the rate of ethanol consumption over time when compared to Cluster 2 (difference = 0.147, *p* = 0.045).

### bQTL Analysis Identifies Significant Locus on Chromosome 14 for Anxiety-Like Behaviors

Significant bQTL (*p* < 0.05) were identified on Chromosome 14 (Chr 14) for transitions between chambers, percent distance traveled in the light, and percent time spent in the light (Figure 3). Transitions between chambers had the highest LOD score (11.59) and most significant *p*-value (< 0.001) of the three phenotypes (Figure 3A), though all three 95% Bayesian C.I.s overlap each other. Founder strain effects are also similar, with the most prominent contribution coming from C57BL/6J alleles which indicated higher levels of anxiety-like behavior. CAST/EiJ alleles across these C.I.s appeared to confer lower levels of anxiety-like behavior. The similarities in position, C.I.s, and haplotype analyses indicate that all three bQTL are influenced by the same genetic factors.

**Figure 3.**
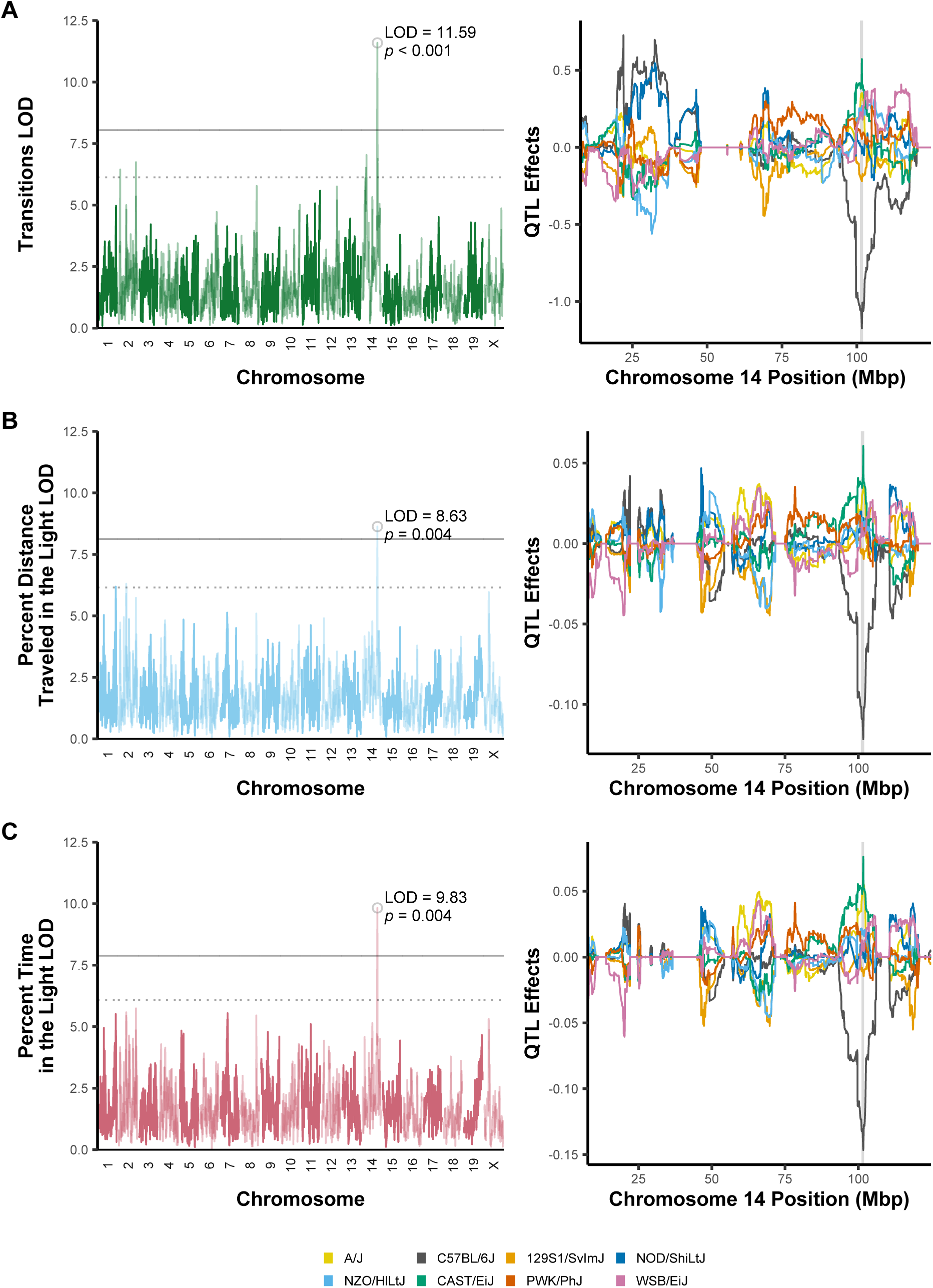
Analysis of anxiety-like behaviors identifies significant QTL on mouse Chromosome 4 with similar founder strain contributions. QTL analysis of transitions between light and dark chambers (A), percent distance traveled in the light chamber (B), and percent time spent in the light chamber (C) all identified a significant locus on Chromosome 14. For all three haplotype analyses, mice with C57BL/6J alleles (black) in the 95% C.I. (shaded region) contributed to decreases in the observed phenotype, indicating higher levels of anxiety-like behavior whereas mice with CAST/EiJ alleles (green) exhibited the opposite effect.

An additional 12 suggestive QTL (*p* < 0.63) and C.I.s were also identified (Table 2), with one for total distance traveled (a measure of locomotor activity) also overlapping the significant QTL on Chr 14. Some of these bQTL overlapped previously-published anxiety-like or locomotor bQTL, most notably the suggestive QTL on Chr 1 for percent distance traveled in the light, which replicated numerous QTL including anxiety-like behaviors from elevated plus maze and light-dark box. [43–47]

**Table 2.** Significant and suggestive QTLs from light-dark box behaviors. QTLs were identified from square root-transformed light-dark box phenotypes and 95% Bayesian C.I.s were calculated using R/qtl2 software. Empirical *p*-values were determined by permutation analysis with 1000 permutations. Previously-published QTLs identified from MGI relate to either anxiety-like behaviors or locomotor activity (for distance traveled and jump count QTLs). QTLs are ordered by *p*-value.

| Phenotype | Chromosome | Peak Position (Mbp) | 95% C.I. (Mbp) | LOD | *p*-value | Percent Variance Explained | Previously Published QTL |
| --- | --- | --- | --- | --- | --- | --- | --- |
| Transitions | 14 | 101.70 | 101.09, 102.08 | 11.59 | 0.001 | 9.79% |  |
| Percent Time in the Light | 14 | 101.68 | 101.09, 101.95 | 9.83 | 0.004 | 8.37% |  |
| Percent Distance Traveled in the Light | 14 | 101.68 | 100.86, 101.98 | 8.63 | 0.004 | 7.39% |  |
| Percent Distance Traveled in the Light | 2 | 75.15 | 15.79, 169.34 | 6.31 | 0.321 | 5.45% | [46] |
| Transitions | 2 | 169.29 | 14.14, 170.07 | 6.74 | 0.327 | 5.82% | [46] |
| Percent Distance Traveled in the Light | 1 | 182.25 | 58.28, 185.97 | 6.19 | 0.378 | 5.35% | [43-47] |
| Distance Traveled | 12 | 97.63 | 71.45, 99.42 | 6.15 | 0.394 | 5.32% |  |
| Distance Traveled | 14 | 105.83 | 68.04, 112.9 | 6.04 | 0.459 | 5.23% |  |
| Distance Traveled | 15 | 93.22 | 92.64, 93.84 | 5.91 | 0.545 | 5.11% |  |
| Distance Traveled | 9 | 106.67 | 32.44, 111.44 | 5.88 | 0.559 | 5.09% | [59] |
| Jump Count | 17 | 66.43 | 27.97, 76.36 | 5.89 | 0.593 | 5.10% | [45, 46, 60] |
| Jump Count | 2 | 141.69 | 44.52, 173.82 | 5.87 | 0.598 | 5.09% | [59] |

### Variant Associations Indicate Potential Role for Gene Expression in Chr 14 QTL

Because transitions between light and dark chambers had the highest LOD score for the Chr 14 QTL, variant associations for that phenotype were estimated and significantly-associated variants were identified using permutation testing across Chr 14 (Figure 4A). Of the 18 significant variants, none fell within the coding region of a gene (Figure 4B); most were intergenic variants, with one additional structural variant (a 391 bp deletion) and one downstream variant for gene *Tbc1d4* (Table 3). Similarly, variant associations were identified for the other two overlapping bQTL on Chr 14 (Figure S5; Figure S6), which additionally contained upstream variants for predicted gene *Gm54405* (Table S2, Table S3). The lack of coding sequence variants indicates that the genetic variants contributing to the significant bQTL on Chr 14 are likely to influence gene expression instead of protein structure.

**Figure 4.**
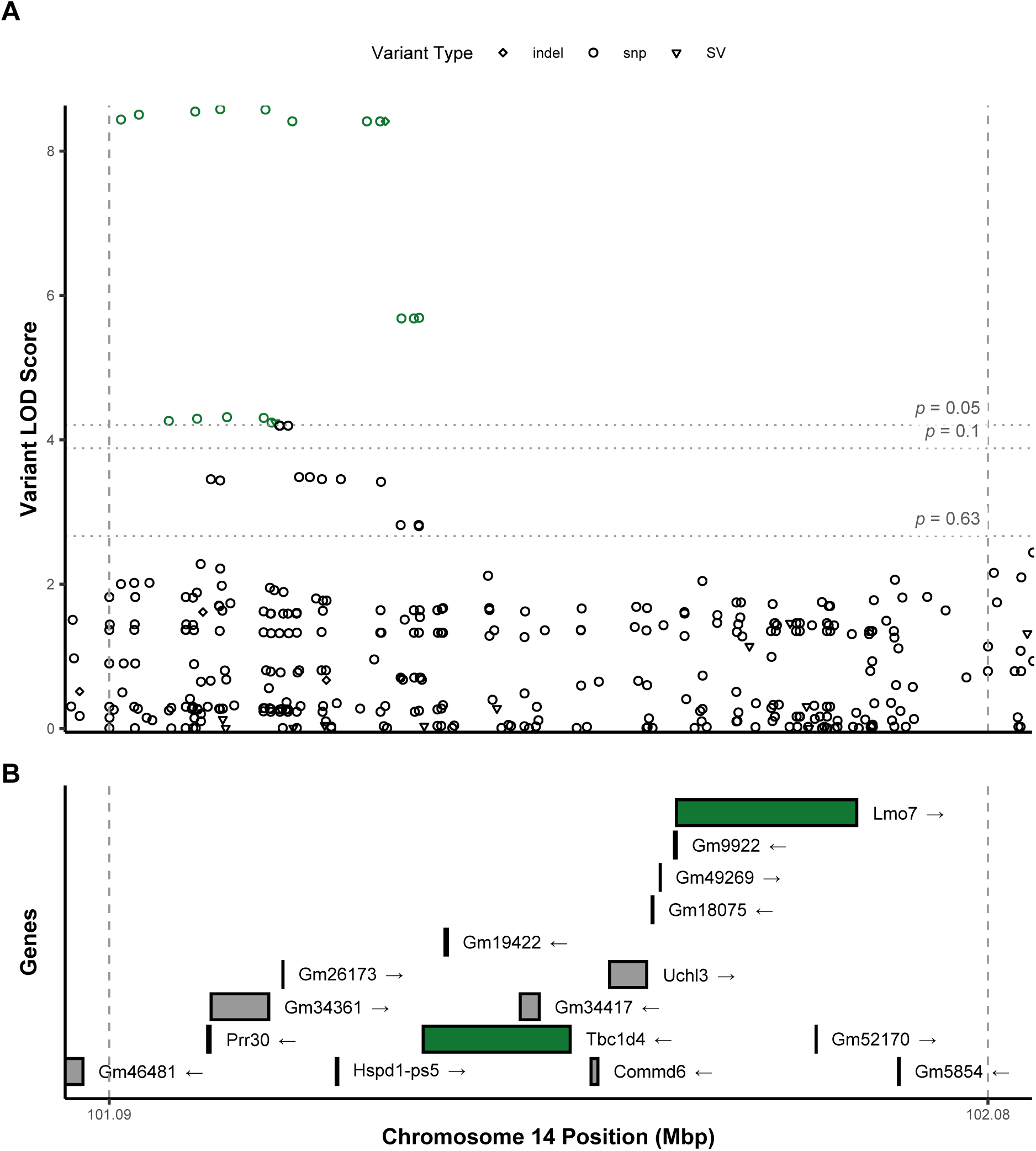
Variant associations across significant bQTL on Chromosome 14 for transitions between light and dark chambers. Associations were estimated across the 95% Bayesian confidence interval identified for the significant bQTL, identified by dashed vertical lines (A). Statistical thresholds estimated by permutation analysis are indicated by horizontal lines; variants with LOD scores surpassing the *p* < 0.05 threshold are highlighted in green. Known gene and pseudogene transcripts identified from MGI databases include *Lmo7* and *Tcb1d4* (B).

**Table 3.**
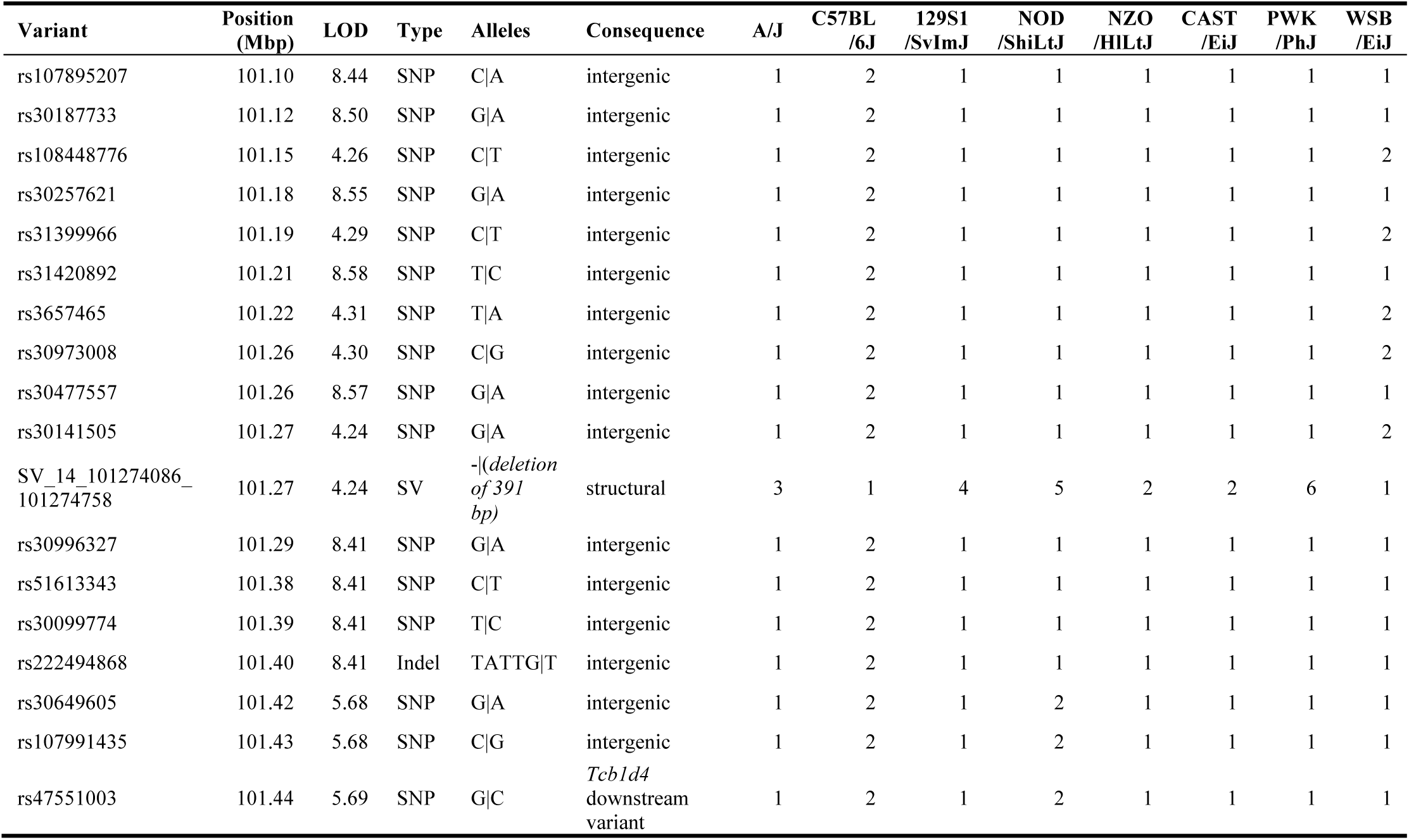
Significant variant associations and strain distributions on Chromosome 14 within QTL 95% C.I. for transitions between light and dark chambers. Variant association LOD scores were calculated using R/qtl2; permutation analysis of 1000 permutations across Chromsome 14 was used to determine empirical significance thresholds. Variant data including consequence and strain distribution pattern were downloaded from MGI. All of the significant variants associated with the phenotype are either intergenic (including one structural variant) except for rs47551003. Variants are ordered by chromosomal location in Mbp.

### eQTL and Bioinformatics Analysis Identify Two Candidate Genes for Anxiety-Like Behavior

RNA-seq data from PFC were used to identify *cis*-eQTL for all 16 genes located within the Chr 14 C.I. for significant bQTL (Table 4). Of these, only one gene had a significant *cis*-eQTL, *Tbc1d4* (Figure 5A). Haplotype analysis shows that mice with PWK/PhJ alleles at this locus had lower levels of *Tbc1d4* expression in PFC, whereas mice with NZO/HILtJ alleles had higher expression (Figure 5B). Two additional genes, *Lmo7* (Figure S7) and *Commd6* (Figure S8), had suggestive *cis*-eQTL.

**Figure 5.**
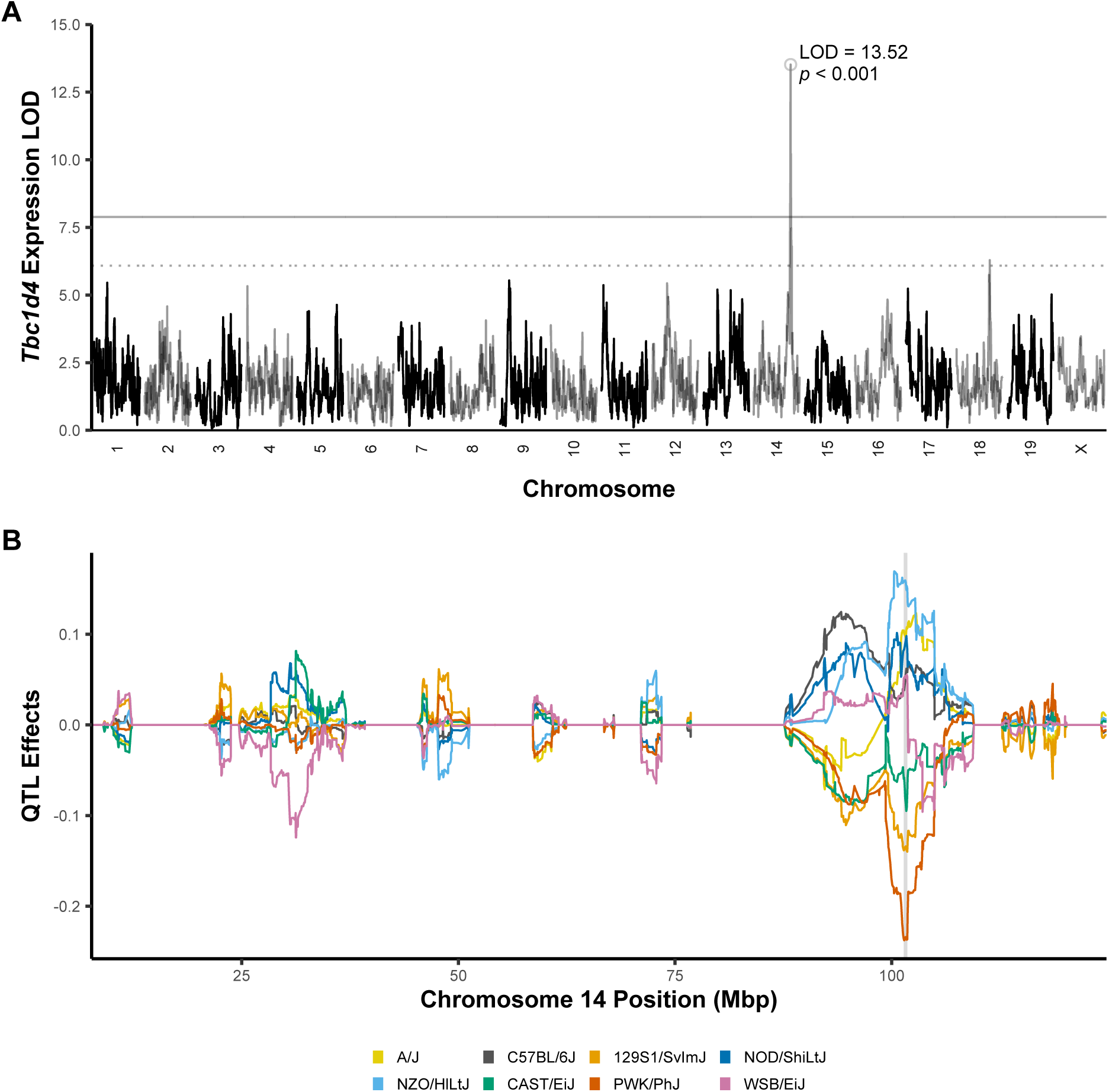
Candidate gene *Tbc1d4* has a significant *cis*-eQTL in PFC overlapping the significant bQTL interval for transitions between light and dark chambers. eQTL analysis of positional candidate genes from bQTL identified *Tbc1d4* as the only gene from the Chr 14 C.I. with a significant *cis*-eQTL in PFC (A). Haplotype analysis shows contributions of PWK/PhJ alleles (red) to lower expression levels and NZO/HILTJ alleles (light blue) to higher expression levels of this gene (B).

**Table 4.** Significant and suggestive *cis*-eQTL from PFC overlapping Chr 14 bQTL C.I. . Of 16 genes and predicted genes in the Chr 14 bQTL 95% Bayesian C.I. for transitions between light and dark chambers, only 3 had *cis*-eQTL that surpassed the *p* < 0.63 threshold. Of these, only *Tbc1d4* had a significant eQTL.

| **MGI Symbol** | **Chromosome** | **Peak Position (Mbp)** | **LOD** | **95% C.I. (Mbp)** | ***p*-value** |
| --- | --- | --- | --- | --- | --- |
| *Tbc1d4* | 14 | 101.74 | 13.52 | 101.39, 101.8 | < 0.001 |
| *Lmo7* | 14 | 100.60 | 7.61 | 28.93, 101.98 | 0.385 |
| *Commd6* | 14 | 101.74 | 6.12 | 99.97, 102.83 | 0.626 |

GWAS Catalog was queried for each gene within the significant bQTL C.I. to find associations with mental health phenotypes. Only one gene, *Lmo7*, had any associations with mental health phenotypes, including obsessive-compulsive disorder [48] and initial pursuit acceleration, a trait associated with psychotic disorders including schizophrenia [49].

## Discussion

### bQTL Analysis Identifies Specific, Novel Loci and Candidate Genes for Anxiety-Like Behaviors

This report is the first to analyze genetic components related to anxiety-like behavior from LDB in DO mice. We identified 3 significant loci on Chr 14 for anxiety-like behaviors including transitions between dark and light chambers, percent distance traveled in the light chamber, and percent time spent in the light chamber (Table 2), each with overlapping C.I.s and similar haplotype effects (Figure 3), which suggests that all 3 phenotypes are influenced by a single locus; this is concordant with the findings that the three phenotypes are significantly correlated (Figure S1). Two high-priority candidate genes were identified from this novel locus, *Tbc1d4* and *Lmo7*.

*Tbc1d4* (TBC1 domain family member 4) is a part of the insulin signaling pathway [50] and is ubiquitously expressed throughout the brain and other tissues [51]. While it has not been previously implicated in anxiety disorders or other psychiatric phenotypes, the presence of a downstream variant with a significant association with the gene (Table 3) and a significant *cis*-eQTL (Table 4; Figure 5) suggests that the same variants associated with anxiety-like behaviors in DO mice are also regulating expression of *Tbc1d4*. The haplotype analyses of anxiety-like behaviors at this locus are mostly driven by C57BL/6J alleles (Figure 3), which are middling at this locus in the eQTL haplotype analysis (Figure 5B); founder strain distribution patterns differ between PFC *Tbc1d4* expression and bQTL phenotypes, but analysis in other brain regions may reveal different founder strain effects. *Tbc1d4* does not appear in any GeneWeaver data sets or human GWAS related to anxiety-like behavior or other psychiatric traits.

*Lmo7* (LIM domain only 7) is also ubiquitously expressed in the brain [51, 52] and is predicted to be involved in transcriptional regulation by MGI. It appears in two data sets from GWAS Catalog [42], where it has a suggestive association with obsessive compulsive disorder (*p* = 3 × 10^−6^) [48] and a significant association with initial pursuit acceleration in psychotic disorders (*p* = 4 × 10^−8^) [49], a trait associated with schizophrenia, bipolar disorder, and schizoaffective disorder. It also has a suggestive *cis*-eQTL (*p* = 0.385) overlapping the significant bQTL interval on Chr 14.

*Lmo7* appears in several datasets related to psychiatric and substance use conditions identified from GeneWeaver. It was differentially expressed in mice following overexpression of *MeCP2*, a gene associated with anxiety disorders [53]. *Lmo7* is also differentially expressed in postmortem human central Amygdala in persons affected by opioid use disorder [54] and dysregulated in dorsolateral PFC of persons with a history of cocaine use [55]. It also has altered expression levels in the PFCs of persons living with AUD [56]. Mouse studies have found it correlates with ethanol withdrawal severity [57] and open field locomotion following cocaine use [24] and that it is differentially expressed in mouse lines bred for high and low acute functional tolerance to ethanol [58]. These findings support *Lmo7* as a putative candidate gene for anxiety-like behaviors.

While the significant bQTL and candidate genes are novel for anxiety-like behaviors, some of the suggestive bQTL overlap with earlier anxiety-like bQTL in the literature (Table 2). bQTL for anxiety-like behaviors in mice on Chr 1, 2, 9, and 17 were found in MGI databases [39] and included behaviors from LDB [43] and other assays such as the elevated plus maze [45] and conditioned fear response [46]. The existence of these mouse bQTL associated with anxiety-like behaviors provides consilience with our results.

### Anxiety-Like Behaviors Predict Voluntary Ethanol Consumption in DO Mice

All anxiety-like behavioral phenotypes from LDB in our study were found to be significantly predictive for voluntary ethanol consumption over the course of four weeks of IEA (Figure 2). While they generally explain only a small amount of the total variance in ethanol consumption, they all have positive coefficient estimates indicating that animals with higher levels of locomotor activity (Figure 2A) and lower levels of anxiety-like behavior (Figure 2B-F) consumed more ethanol over the course of IEA.

This finding is consistent with the results of *k*-means clustering by LDB phenotypes (Figure S2), where Cluster 1 exhibited lower levels of anxiety-like behavior but consumed significantly more ethanol over the course of the study (Figure S3A). Strikingly, cluster means differed significantly even in week four of IEA and held to the same pattern even though clusters were based on LDB assays run prior to initiation of IEA, demonstrating some stability in phenotypes.

### Limitations

Despite the novel findings of this report, several limitations should be considered. First, the DO mouse population included in this study only consisted of male animals due to power and space considerations. Future studies are planned to validate identified loci in female mice, but the current findings may not be generalizable to females. Second, RNA-seq data used for these analyses was collected from bulk tissue samples in PFC and the majority of samples came from animals following ethanol exposure. However, *cis*-eQTL are associated with higher heritability than *trans*-eQTL [51] and here are assumed to be the result of genetic factors rather than modulated by ethanol exposure. Future studies may focus on ethanol-naïve animals, single-cell RNA-seq studies, and studies of brain regions such as the amygdala that have been implicated in anxiety disorders.

### Conclusions

These findings identified novel candidate genes *Tbc1d4* and Lmo7 and loci associated with anxiety-like behaviors, adding to our understanding of the complex genetic architecture of anxiety disorders. Further transcriptomic, bioinformatic, and behavioral studies involving these candidate genes promise to reveal mechanistic and causal insights into these findings. Importantly, this work also identified basal anxiety genetic subgroups that are predictive of later ethanol consumption phenotypes. This finding could have important implications on factors influencing risk for progression to alcohol use disorder in humans.

## Acronyms

DO: Diversity Outbred
GWAS: Genome-Wide Association Study
IEA: Intermittent Ethanol Access
LDB: Light-Dark Box
LOD: Logarithm of the Odds
PFC: Prefrontal Cortex
QTL: Quantitative Trait Locus / Loci
bQTL: Behavioral Quantitative Trait Locus / Loci
eQTL: Expression Quantitative Trait Locus / Loci

---

**Supplemental Table 1.**
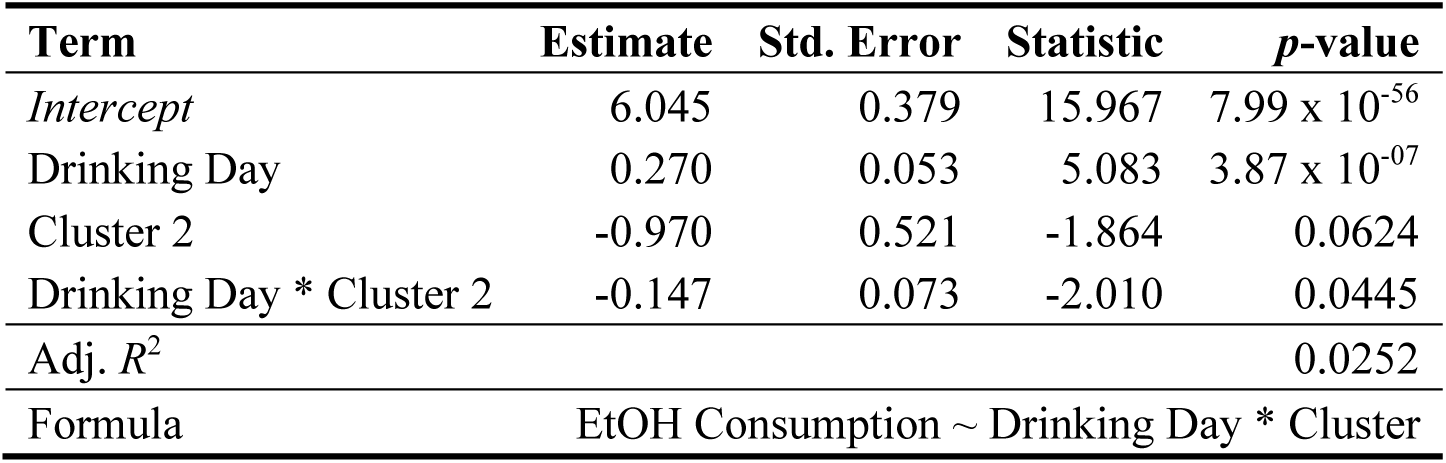
Trajectory of ethanol consumption over time differs across *k*-means clusters generated from anxiety-like behaviors. Mice were grouped into two clusters based on LDB phenotypes. A multiple linear model was fitted to predict ethanol consumption using drinking day, cluster, and an interaction term. The significant interaction term (*p* = 0.045) indicates that the slopes significantly differ between clusters, with Cluster 2 showing less of a progressive increase in ethanol consumption over time compared to Cluster 1.

**Supplemental Table 2.** Significant variant associations and strain distributions on Chromosome 14 within QTL 95% C.I. for percent distance traveled in the light chamber. Variant association LOD scores were calculated using R/qtl2; permutation analysis of 1000 permutations across Chromsome 14 was used to determine empirical significance thresholds. Variant data including consequence and strain distribution pattern were downloaded from MGI. All of the significant variants associated with the phenotype are either intergenic (including one structural variant) except for rs50653690 and rs31416866, which are upstream variants for predicted gene *Gm54405*. Variants are ordered by chromosomal location in Mbp.

| Variant | Position (Mbp) | LOD | Type | Alleles | Consequence | A/J | C57BL /6J | 129S1 /SvImJ | NOD /ShiLtJ | NZO /HILtJ | CAST /EiJ | PWK /PhJ | WSB /EiJ |
| --- | --- | --- | --- | --- | --- | --- | --- | --- | --- | --- | --- | --- | --- |
| rs31086859 | 100.89 | 6.24 | SNP | G\|A | intergenic | 1 | 2 | 1 | 1 | 1 | 1 | 1 | 1 |
| rs31397583 | 100.90 | 6.24 | SNP | G\|A | intergenic | 1 | 2 | 1 | 1 | 1 | 1 | 1 | 1 |
| rs30239536 | 100.97 | 6.23 | SNP | G\|A | intergenic | 1 | 2 | 1 | 1 | 1 | 1 | 1 | 1 |
| rs30556413 | 101.02 | 6.58 | SNP | G\|A | intergenic | 1 | 2 | 1 | 1 | 1 | 1 | 1 | 1 |
| rs107895207 | 101.10 | 6.97 | SNP | C\|A | intergenic | 1 | 2 | 1 | 1 | 1 | 1 | 1 | 1 |
| rs30187733 | 101.12 | 7.06 | SNP | G\|A | intergenic | 1 | 1 | 1 | 1 | 1 | 1 | 1 | 1 |
| rs108448776 | 101.15 | 4.11 | SNP | C\|T | intergenic | 1 | 2 | 1 | 1 | 1 | 1 | 1 | 2 |
| rs30257621 | 101.18 | 7.09 | SNP | G\|A | intergenic | 1 | 2 | 1 | 1 | 1 | 1 | 1 | 1 |
| rs31399966 | 101.19 | 4.13 | SNP | C\|T | intergenic | 1 | 2 | 1 | 1 | 1 | 1 | 1 | 2 |
| rs31420892 | 101.21 | 7.12 | SNP | T\|C | intergenic | 1 | 2 | 1 | 1 | 1 | 1 | 1 | 1 |
| rs3657465 | 101.22 | 4.15 | SNP | T\|A | intergenic | 1 | 2 | 1 | 1 | 1 | 1 | 1 | 2 |
| rs30973008 | 101.26 | 4.16 | SNP | C\|G | intergenic | 1 | 2 | 1 | 1 | 1 | 1 | 1 | 2 |
| rs30477557 | 101.26 | 7.14 | SNP | G\|A | intergenic | 1 | 2 | 1 | 1 | 1 | 1 | 1 | 1 |
| rs30141505 | 101.27 | 4.14 | SNP | G\|A | intergenic | 1 | 2 | 1 | 1 | 1 | 1 | 1 | 2 |
| SV_14_101274086_101274758 | 101.27 | 4.14 | SV | -(deletion of 391 bp) | structural | 3 | 1 | 4 | 5 | 2 | 2 | 6 | 1 |
| rs50653690 | 101.28 | 4.12 | SNP | T\|C | *Gm54405* upstream variant | 1 | 2 | 1 | 1 | 1 | 1 | 1 | 2 |
| rs31416866 | 101.28 | 4.12 | SNP | T\|A | *Gm54405* upstream variant | 1 | 2 | 1 | 1 | 1 | 1 | 1 | 2 |
| rs30979977 | 101.29 | 4.12 | SNP | T\|C | intergenic | 1 | 2 | 1 | 1 | 1 | 1 | 1 | 2 |
| rs30996327 | 101.29 | 7.10 | SNP | G\|A | intergenic | 1 | 2 | 1 | 1 | 1 | 1 | 1 | 1 |
| rs51613343 | 101.38 | 7.10 | SNP | C\|T | intergenic | 1 | 2 | 1 | 1 | 1 | 1 | 1 | 1 |
| rs30099774 | 101.39 | 7.10 | SNP | T\|C | intergenic | 1 | 2 | 1 | 1 | 1 | 1 | 1 | 1 |
| rs222494868 | 101.40 | 7.10 | indel | TATTG\|T | intergenic | 1 | 2 | 1 | 1 | 1 | 1 | 1 | 1 |

**Supplemental Table 3.** Significant variant associations and strain distributions on Chromosome 14 within QTL 95% C.I. for percent time spent in the light chamber. Variant association LOD scores were calculated using R/qtl2; permutation analysis of 1000 permutations across Chromsome 14 was used to determine empirical significance thresholds. Variant data including consequence and strain distribution pattern were downloaded from MGI. All of the significant variants associated with the phenotype are either intergenic (including one structural variant) except for rs50x653690, rs31416866, and rs47551003. Variants are ordered by chromosomal location in Mbp.

| Variant | Position (Mbp) | LOD | Type | Alleles | Consequence | A/J | C57BL /6J | 129S1 /SvImJ | NOD /ShiLtJ | NZO /HILtJ | CAST /EiJ | PWK /PhJ | WSB /EiJ |
| --- | --- | --- | --- | --- | --- | --- | --- | --- | --- | --- | --- | --- | --- |
| rs107895207 | 101.10 | 7.52 | SNP | C\|A | intergenic | 1 | 2 | 1 | 1 | 1 | 1 | 1 | 1 |
| rs30187733 | 101.12 | 7.61 | SNP | G\|A | intergenic | 1 | 2 | 1 | 1 | 1 | 1 | 1 | 1 |
| rs108448776 | 101.15 | 4.59 | SNP | C\|T | intergenic | 1 | 2 | 1 | 1 | 1 | 1 | 1 | 2 |
| rs30257621 | 101.18 | 7.63 | SNP | G\|A | intergenic | 1 | 2 | 1 | 1 | 1 | 1 | 1 | 1 |
| rs31399966 | 101.19 | 4.61 | SNP | C\|T | intergenic | 1 | 2 | 1 | 1 | 1 | 1 | 1 | 2 |
| rs31420892 | 101.21 | 7.66 | SNP | T\|C | intergenic | 1 | 2 | 1 | 1 | 1 | 1 | 1 | 1 |
| rs3657465 | 101.22 | 4.63 | SNP | T\|A | intergenic | 1 | 2 | 1 | 1 | 1 | 1 | 1 | 2 |
| rs30973008 | 101.26 | 4.62 | SNP | C\|G | intergenic | 1 | 2 | 1 | 1 | 1 | 1 | 1 | 2 |
| rs30477557 | 101.26 | 7.64 | SNP | G\|A | intergenic | 1 | 2 | 1 | 1 | 1 | 1 | 1 | 1 |
| rs30141505 | 101.27 | 4.54 | SNP | G\|A | intergenic | 1 | 2 | 1 | 1 | 1 | 1 | 1 | 2 |
| SV_14_101274086_101274758 | 101.27 | 4.54 | SV | -(deletion of 391 bp) | structural | 3 | 1 | 4 | 5 | 2 | 2 | 6 | 1 |
| rs50653690 | 101.28 | 4.50 | SNP | T\|C | *Gm54405* upstream variant | 1 | 2 | 1 | 1 | 1 | 1 | 1 | 2 |
| rs31416866 | 101.28 | 4.50 | SNP | T\|A | *Gm54405* upstream variant | 1 | 2 | 1 | 1 | 1 | 1 | 1 | 2 |
| rs30979977 | 101.29 | 4.50 | SNP | T\|C | intergenic | 1 | 2 | 1 | 1 | 1 | 1 | 1 | 2 |
| rs30996327 | 101.29 | 7.48 | SNP | G\|A | intergenic | 1 | 2 | 1 | 1 | 1 | 1 | 1 | 1 |
| rs51613343 | 101.38 | 7.47 | SNP | C\|T | intergenic | 1 | 2 | 1 | 1 | 1 | 1 | 1 | 1 |
| rs30099774 | 101.39 | 7.47 | SNP | T\|C | intergenic | 1 | 2 | 1 | 1 | 1 | 1 | 1 | 1 |
| rs222494868 | 101.40 | 7.47 | Indel | TATTG\|T | intergenic | 1 | 2 | 1 | 1 | 1 | 1 | 1 | 1 |
| rs30649605 | 101.42 | 4.80 | SNP | G\|A | intergenic | 1 | 2 | 1 | 2 | 1 | 1 | 1 | 1 |
| rs107991435 | 101.43 | 4.80 | SNP | C\|G | intergenic | 1 | 2 | 1 | 2 | 1 | 1 | 1 | 1 |
| rs47551003 | 101.44 | 4.82 | SNP | G\|C | *Tcb1d4* downstream variant | 1 | 2 | 1 | 2 | 1 | 1 | 1 | 1 |

**Supplemental Figure 1.**
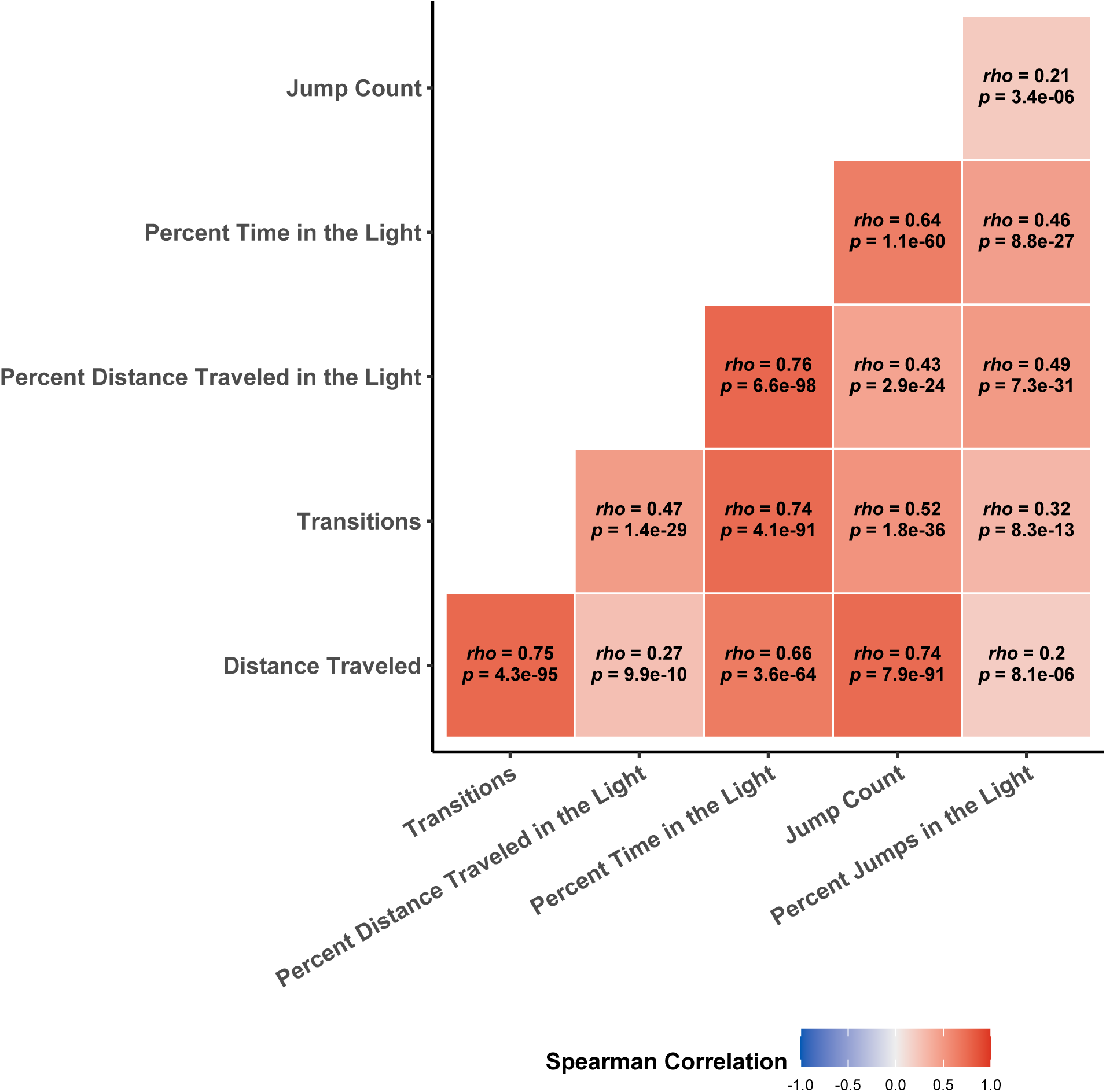
Light-dark box behaviors are significantly correlated. Spearman rank correlations and *p*-values were calculated pairwise for each phenotype. Correlation values range from 0.2 – 0.76 and all are significant following Holm correction.

**Supplemental Figure 2.**
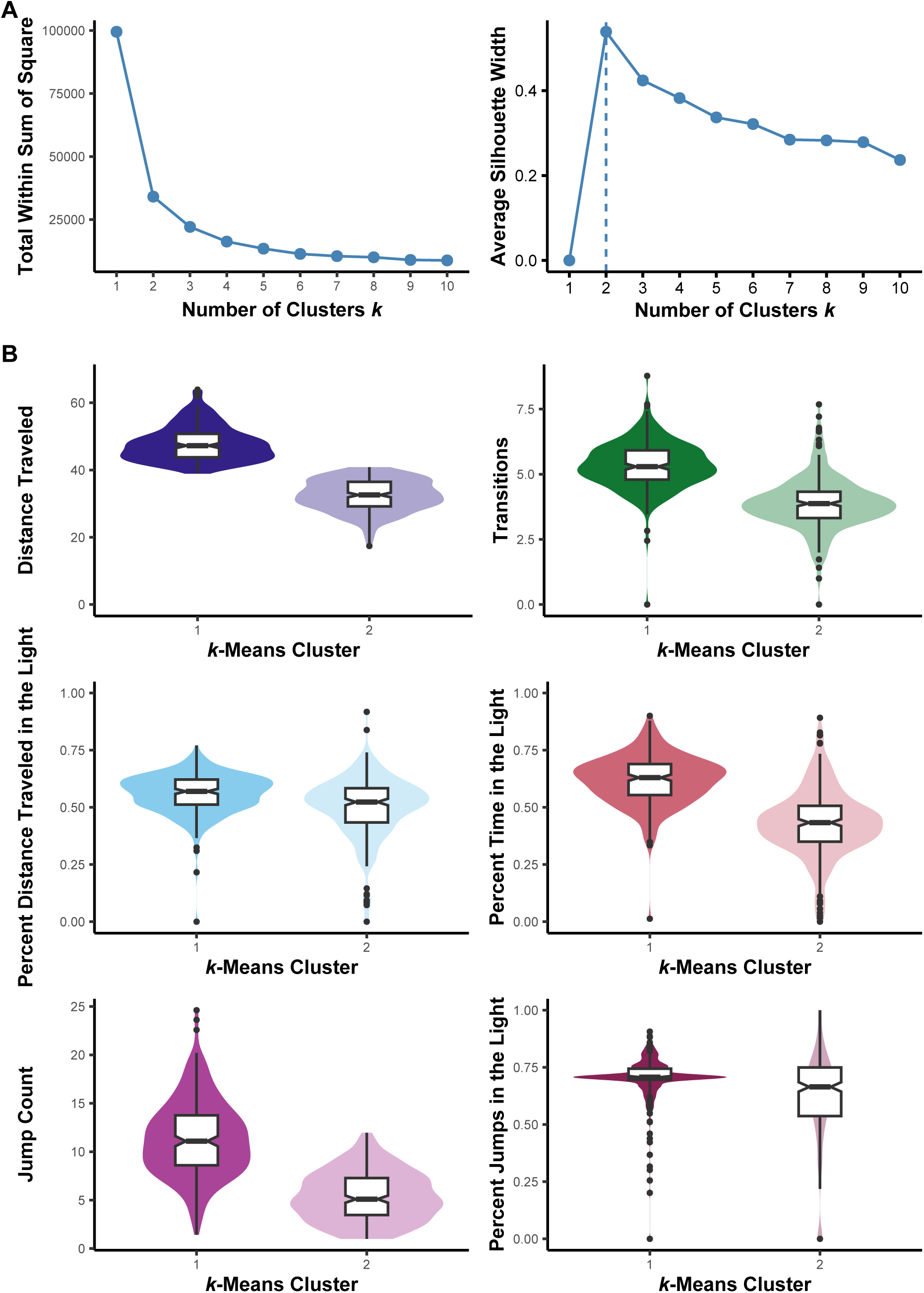
Determination of *k*-means clusters using light-dark box phenotypes. Both the total within sum of squares and average silhouette width methods suggested that two clusters of mice existed within the data set (A). Across phenotypes, Cluster 1 exhibited higher locomotor activity and lower anxiety-like behavior compared to Cluster 2, as evidenced by higher numbers of transitions between chambers, percent distance traveled and percent time spent in the light chamber (B).

**Supplemental Figure 3.**
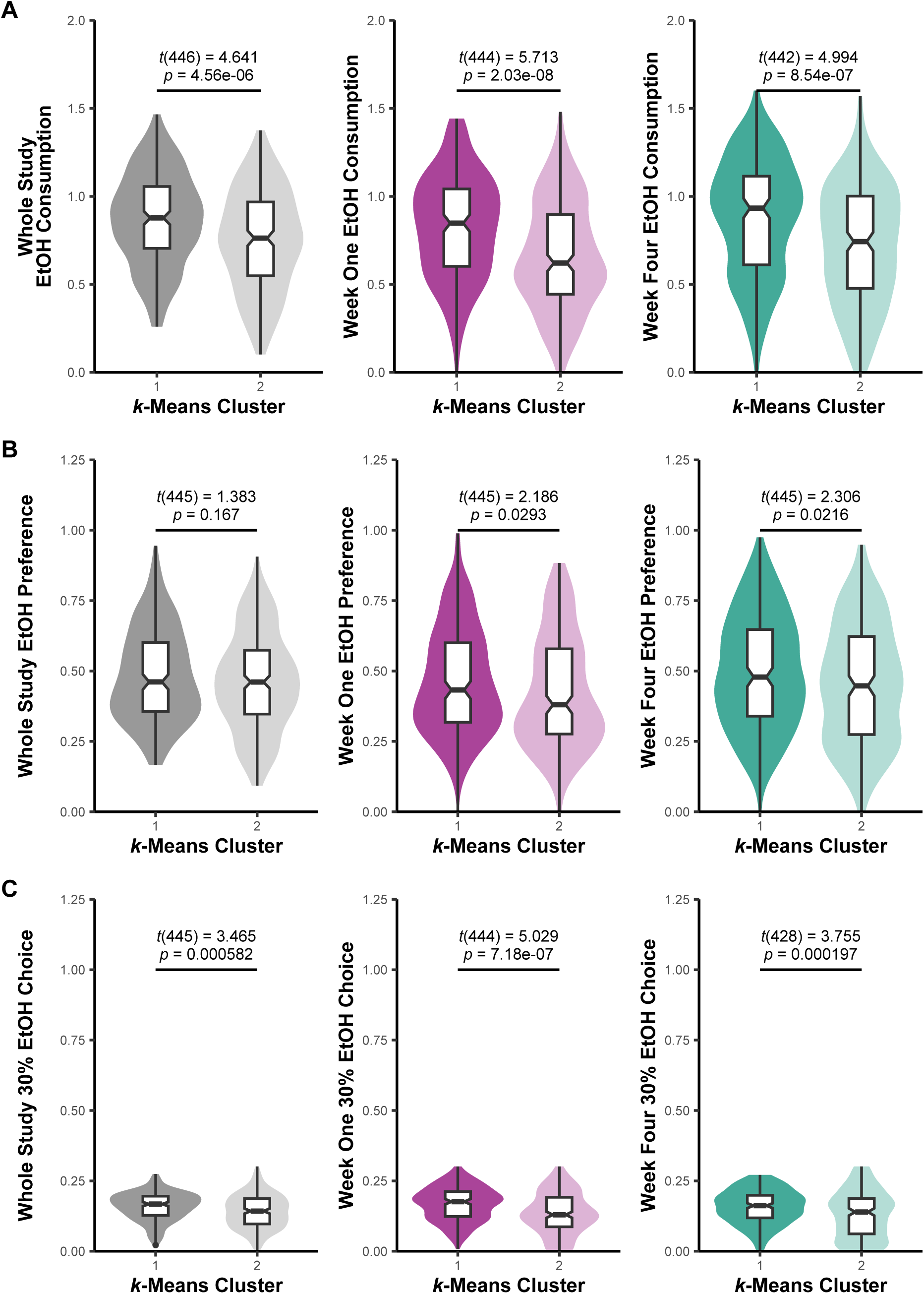
Ethanol consumption behaviors differ across *k*-means clusters generated from anxiety-like behaviors. Following *k*-means clustering of mice into two groups using LDB phenotypes, cluster means for whole study, week one, and week four ethanol consumption (A), preference (B), and 30% ethanol choice (C) were compared using *t*-tests. Ethanol consumption and 30% ethanol choice were log-transformed for normalization; ethanol preference was square-root transformed. Overall, the mean differences appear significant for all comparisons except whole study ethanol preference, with Cluster 1 (associated with higher locomotor activity and lower anxiety-like behavior) having higher ethanol consumption, preference, and 30% ethanol choice than Cluster 2.

**Supplemental Figure 4.**
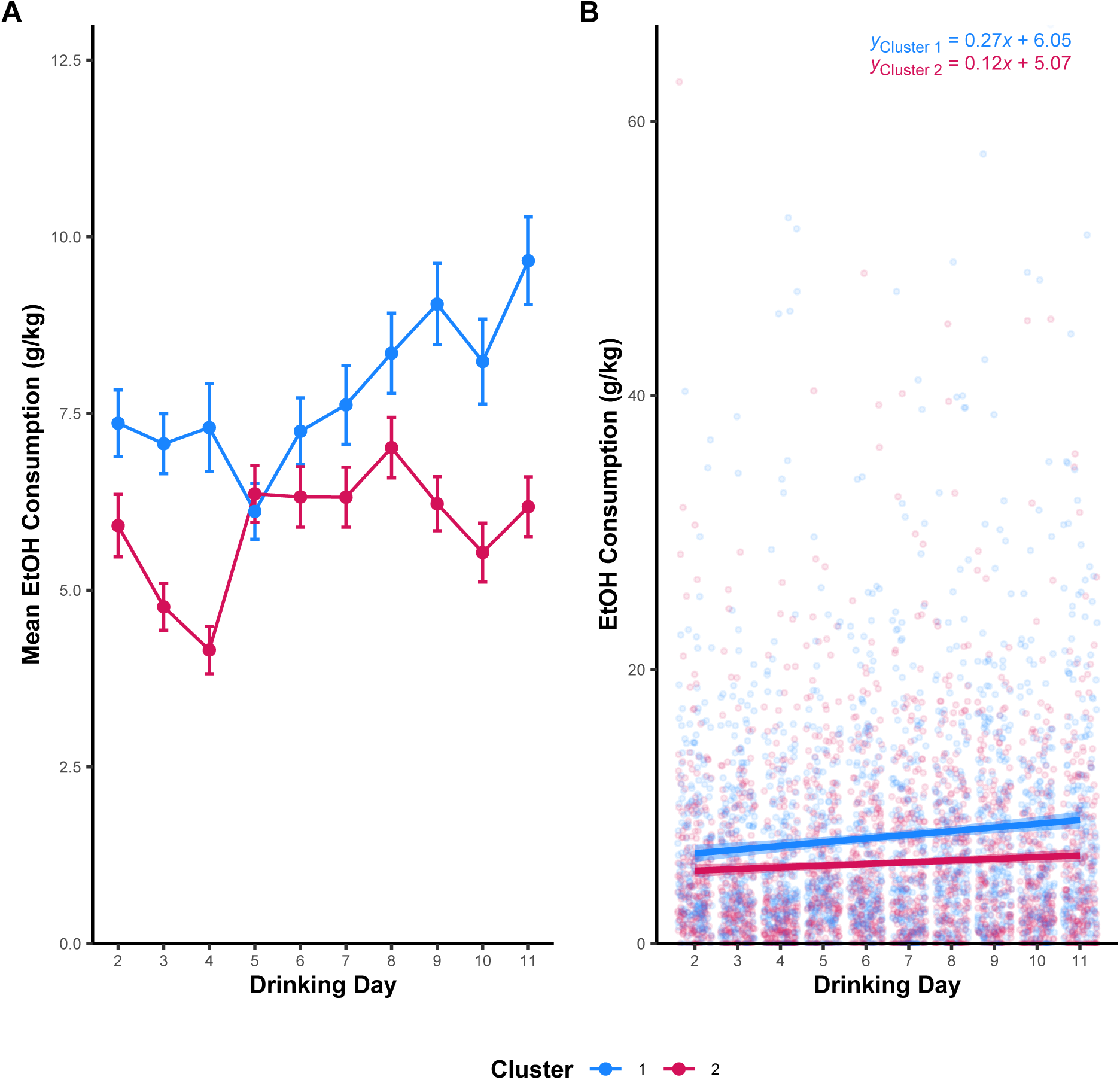
Trajectory of ethanol consumption over time differs across *k*-means clusters generated from anxiety-like behaviors. Following *k*-means clustering of mice into two groups using LDB phenotypes, daily mean ethanol consumption from IEA appeared to differ between the two clusters (A). To test if the rate at which ethanol consumption increased over the course of IEA differed between the clusters, a multiple linear model was fitted with formula ‘EtOH Consumption ∼ Drinking Day * Cluster’ (B). Results indicated a significant difference in the slope of ethanol consumption over time across the two clusters (*p* = 0.045; see Table S1).

**Supplemental Figure 5.**
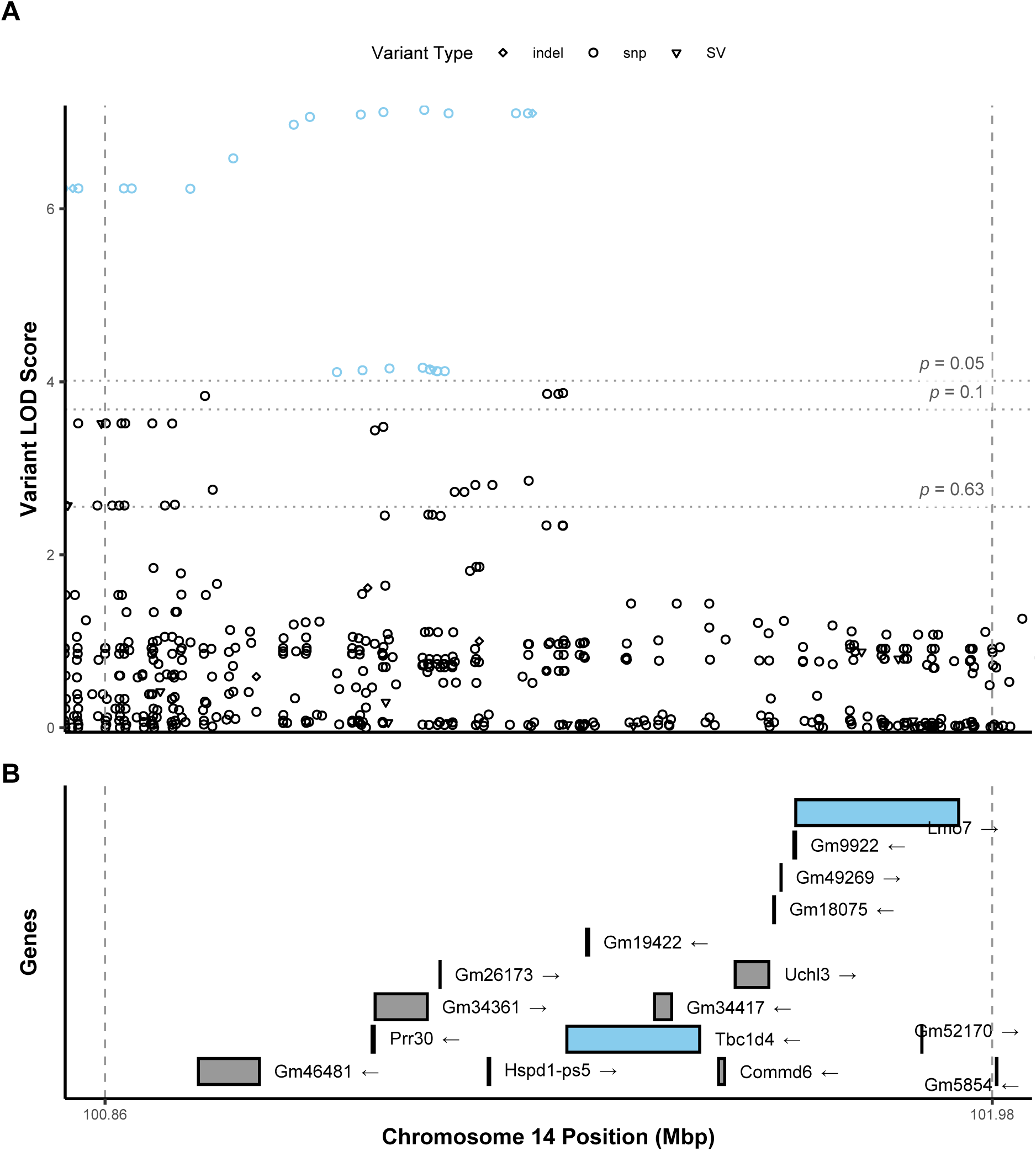
Variant associations across significant bQTL on Chromosome 14 for percent distance traveled in the light chamber. Associations were estimated across the 95% Bayesian confidence interval identified for the significant bQTL, identified by dashed vertical lines (A). Statistical thresholds estimated by permutation analysis are indicated by horizontal lines; variants with LOD scores surpassing the *p* < 0.05 threshold are highlighted in blue. Known gene and pseudogene transcripts identified from MGI databases include *Lmo7* and *Tcb1d4* (B).

**Supplemental Figure 6.**
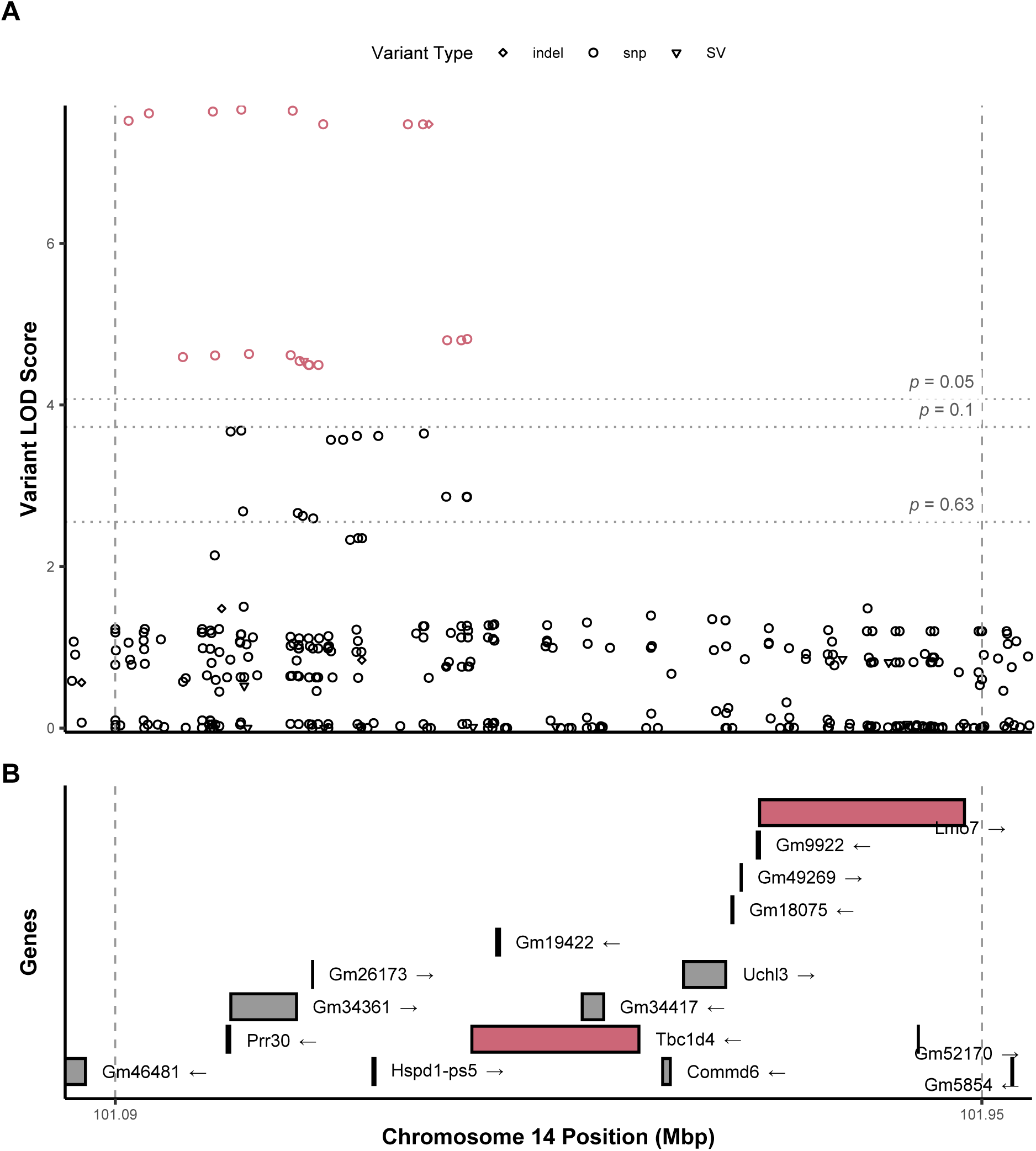
Variant associations across significant bQTL on Chromosome 14 for percent time in the light chamber. Associations were estimated across the 95% Bayesian confidence interval identified for the significant bQTL, identified by dashed vertical lines (A). Statistical thresholds estimated by permutation analysis are indicated by horizontal lines; variants with LOD scores surpassing the *p* < 0.05 threshold are highlighted in red. Known gene and pseudogene transcripts identified from MGI databases include *Lmo7* and *Tcb1d4* (B).

**Supplemental Figure 7.**
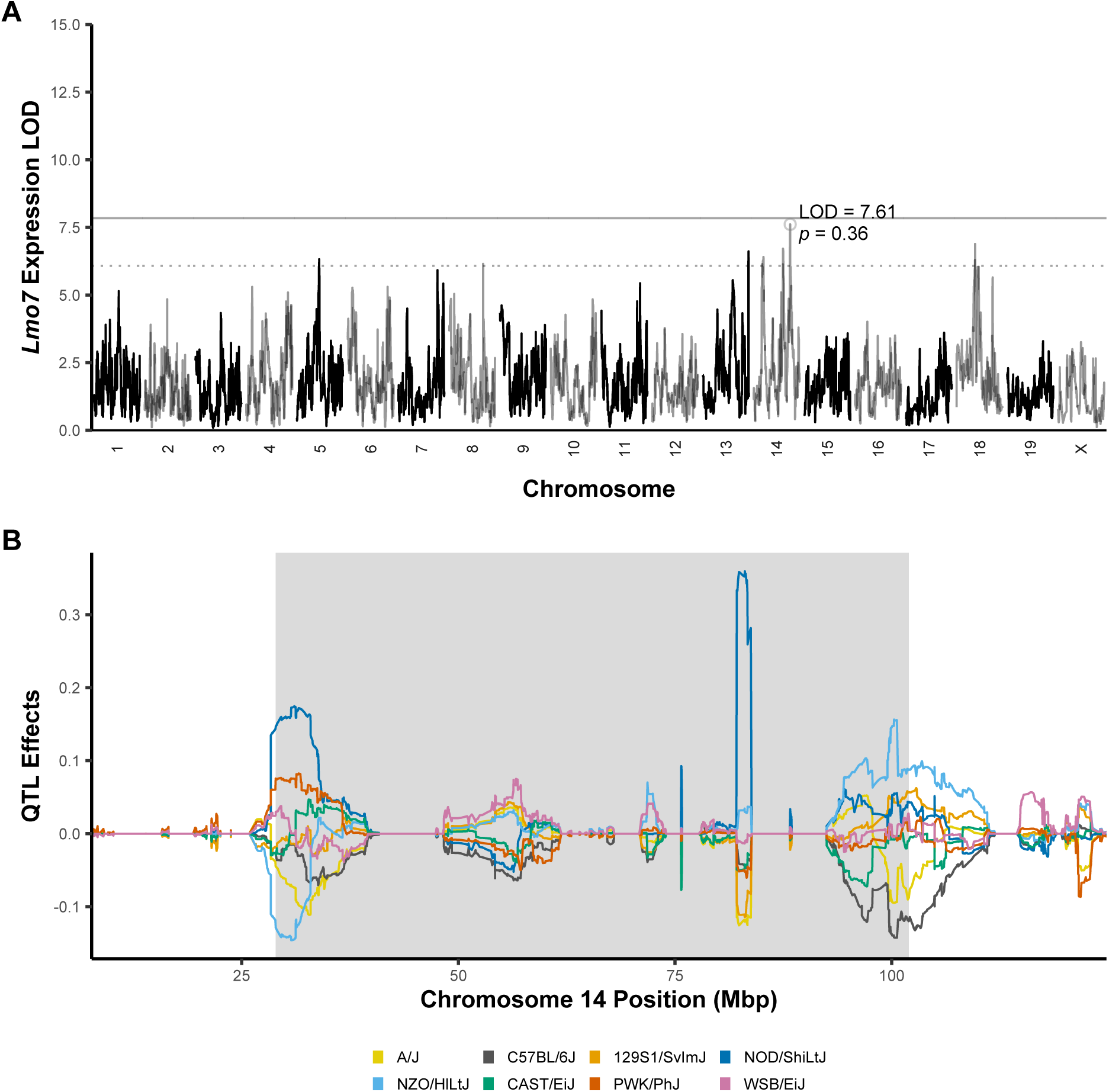
L*m*o7 has a suggestive eQTL in PFC overlapping the significant bQTL interval for transitions between light and dark chambers. eQTL analysis of positional candidate genes from bQTL identified a suggestive *cis*-eQTL for *Lmo7* in PFC (A). Haplotype analysis shows primarily a contribution of NZO/HILtJ alleles (light blue) across the eQTL 95% C.I. (shaded) contributing to increased expression of this gene and C57BL/6J alleles (black) contributing to decreased expression (B).

**Supplemental Figure 8.**
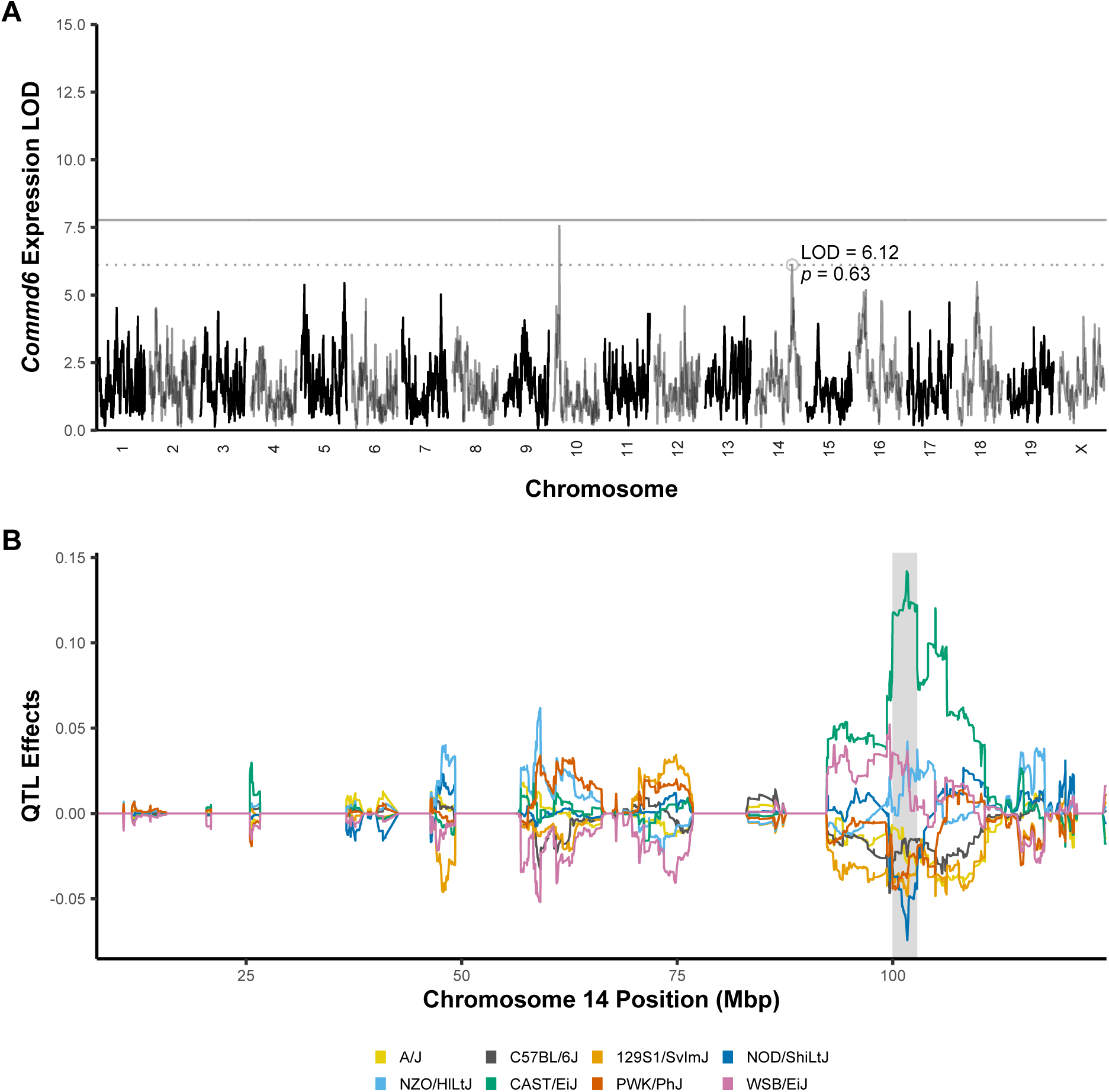
C*o*mmd6 has a suggestive eQTL in PFC overlapping the significant bQTL interval for transitions between light and dark chambers. eQTL analysis of positional candidate genes from bQTL identified a suggestive *cis*-eQTL for *Commd6* in PFC (A). Haplotype analysis shows primarily a contribution of CAST/EiJ alleles (green) across the eQTL 95% C.I. (shaded) contributing to increased expression of this gene (B).

## Notes

### Competing Interest Statement

The authors have declared no competing interest.

### Funding

- National Institute on Alcohol Abuse and Alcoholism: P50AA022537, 1F31AA031189

